# Task engagement differentially drives hippocampal and neocortical neural codes

**DOI:** 10.1101/2025.06.26.661816

**Authors:** Cantin Ortiz, Manuela Allegra, Christoph Schmidt-Hieber

## Abstract

Sensory inputs are progressively transformed into internal representations of the environment along the cortical hierarchy. How does the behavioral relevance of these inputs affect this encoding? Using two-photon calcium imaging in mice navigating virtual-reality environments, we found that visual cortex maintained high sensory discrimination regardless of behavioural engagement, whereas the hippocampal dentate gyrus required active navigation for effective discrimination. These findings suggest that sensory cortices act as general-purpose sensory discriminators, while the hippocampus filters information based on task relevance.

---

The hippocampus is essential for episodic memory and spatial navigation, organizing neuronal activity into a cognitive map of the environment, which emerges from multisensory integration (Burgess et al., 2002; Eichenbaum, 2017; Moser et al., 2017). Sensory information from the visual environment, for instance, is first processed in the primary visual cortex (V1), where neurons encode low-level features like orientation and length of visual stimuli (Lee et al., 1998). However, recent studies have revealed that even at this early stage, V1 responses are modulated by behavior, context and spatial location (Dadarlat and Stryker, 2017; Goltstein et al., 2013; Henschke et al., 2020; Saleem et al., 2018). In parallel, hippocampal spatial codes are highly sensitive to the behavioral state (Allegra et al., 2020; G. Chen et al., 2013; Pettit et al., 2022; Sosa et al., 2025), and remarkably, hippocampal neurons can respond to elementary sensory stimuli—such as a moving bar of light—that are typically associated with processing in primary sensory cortices (Purandare et al., 2022).

While both V1 and hippocampal neurons encode visual sensory inputs, behaviour and space, we hypothesized that sensory processing in the two regions serves distinct functions. As the visual cortex broadcasts information to various brain regions, it needs to produce a faithful representation of the visual environment that gradually changes with variations in visual inputs. In contrast, hippocampal representations converge towards distinct states representing different environments as a function of behavioral relevance. To test this hypothesis, we first established a behavioural task where animals had to discriminate between different visual environments using an immersive virtual reality (VR) setup for head-fixed mice running on a linear treadmill (Fig. 1a) (Allegra et al., 2020). The three visual environments (A, A’ and B) consisted of linear corridors (1.3 m length) with different wall gratings and local cues (Fig. 1b). A and A’ only differed in the orientation of gratings on wall textures, whereas A and B differed considerably in landmarks, textures and colors. In order to receive a drop of sucrose solution, mice were required to stop within an uncued reward zone, which differed in location between environments. After completing a lap, mice were teleported back to the start of either one of the corridors at random. A preliminary training phase comprising five sessions, during which the animals were only exposed to environment A, was sufficient for them to learn the task, as quantified by several behavioural parameters (Fig. S1a,b) such as an increased lick rate (Fig. 1c) and reduced running speed (Fig. 1d) in anticipation of the delivery of the reward. Following the completion of the training phase, the animals were exposed to either a random alternation between the two similar environments A and A’, or between the two more distinct corridors A and B.

**Figure 1.**
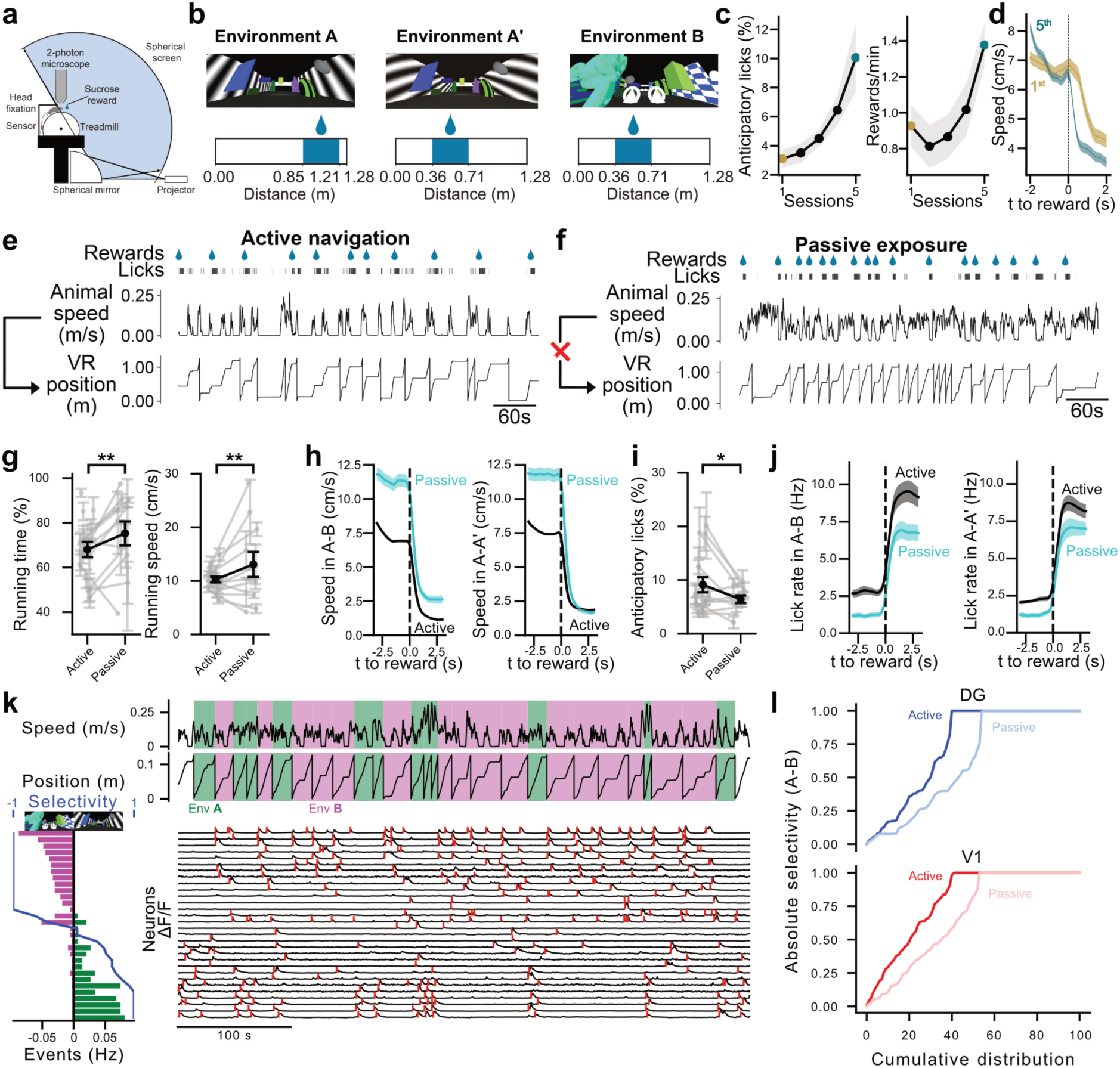
Neuronal discrimination during a visual discrimination task in virtual reality depends on behavioural engagement in both V1 and DG **(a)** Schematic of the behavioural setup. **(b)** View of the three environments used from the starting location. Reward zone is displayed in blue. **(c)** Learning curves of the animals during the first 5 sessions. **(d)** Comparison of speed averaged around reward delivery on day 1 (mustard) and day 5 (cyan). **(e)** Principle of active navigation in VR. **(f)** Principle of passive exposure in VR, with decoupling between animal speed and VR position. **(g)** Quantification of locomotion during active and passive exposure for the environment pair A-B. **(h)** Speed averaged around reward delivery during active (black) and passive (cyan) conditions, in the A-B pair of environments (left) and the A-A’ pair (right). **(i)** Comparison of anticipatory licking during active and passive exposure for the A-B pair of environments. **(j)** Lick rate averaged around reward delivery during active (black) and passive (cyan) conditions, in the A-B pair of environments (left) and the A-A’ pair (right). **(k)** Example of a recording session. Top: running speed and inferred position in the virtual environment as a function of time. Visited environments are indicated by the background colour (green: A, magenta: B). Bottom right: calcium traces from a subset of neurons sorted by increasing selectivity. Identified calcium transients are marked with a red vertical line. Bottom left: mean event rate of each neuron in environments A and B (green and magenta bars). Selectivity value is displayed in blue. **(l)** Distribution of absolute selectivity from individual neurons pulled across A-B recording sessions in active (darker colours) and passive (lighter) conditions, for DG (top) and V1 recordings (bottom). In **(c)**, **(d)**, **(h)**, **(j)**, solid lines show the mean while shaded areas correspond to the standard error of the mean. In **(g)** and **(i)**, error bars indicate the standard error of the mean. Grey lines correspond to multiple sessions from a single animal, while dark lines show the average across animals.

During active navigation, the animal’s position along the corridor was determined by the distance run by the animal on the wheel (Fig. 1e). To assess the effect of behavioural engagement on visual discrimination, a passive exposure paradigm was devised, where sensory inputs presented to the animal were similar to those in the active condition, but uncoupled from the animal’s movement on the treadmill. Instead of converting the running speed into a position within the virtual reality (VR), a replay from another session was displayed (Fig. 1f), thereby entirely decoupling the animal’s locomotion from the displayed environment (Allegra et al., 2020). The rewards were delivered within one of the two reward zones, but the reward zones were randomly selected for each lap crossing independently of the environment that was currently presented. This randomized reward delivery was maintained to selectively abolish the task goal while minimizing other changes to the context. In the passive exposure paradigm, the animals ran longer and faster compared to active navigation (Fig. 1g, Fig. S1c,d). Furthermore, the animals did not display anticipatory deceleration, showed a reduced tendency to stop following the delivery of a reward (Fig. 1h) and to lick in anticipation of a reward in the A-B pair of environments (Fig. 1i, Fig. S1e). The reward-triggered average indicates that the lick rate was increasing prior to reward delivery during active navigation, but not during passive exposure (Fig. 1j). During active navigation, animals performed slightly worse in the A-A’ pair of environments due to a decreased performance in A’ compared to B (Fig. S1f-j). Possibly reflecting this lower performance, they also ran faster in A’ compared to B during active navigation, but not during passive exposure (Fig. S1k-m). Together, the task structure enables us to distinguish between active behavioural engagement on the one hand and passive exposure to a pre-recorded movie without the animal’s active participation on the other hand.

To monitor neuronal population activity during the discrimination task, we performed *in vivo* two-photon calcium imaging from either the hippocampus or the visual cortex of head-fixed mice executing the discrimination task described above. A total of two distinct groups of animals were injected with the genetically encoded calcium indicator GCaMP6f and implanted with optical devices in either the primary visual cortex (V1) or the dentate gyrus (DG) subregion of the hippocampus. Neuronal activity and animal behaviour were recorded over a period of approximately 500 seconds per session. During recordings, some animals were only subject to the active navigation paradigm, while others were recorded during both active navigation and passive exposure. The inferred neuronal activity was derived from calcium traces using a thresholding approach to detect calcium events (Allegra et al., 2020; Dombeck et al., 2010), which are likely to arise from bursts of action potentials (Fig. 1k, right). To assess whether neurons fired more in one environment than in the other one, we first computed the selectivity of individual neurons (Fig. 1k, left). Taking together selectivity values across different sessions revealed that during active navigation, environments seemed to be more accurately discriminated by neuronal activity than during passive exposure in both brain regions, though the difference appeared to be less pronounced in V1 than in the dentate gyrus (Fig. 1l, Fig. S1m).

To further explore these differences, we first sought to better understand how our experimental variables affected the rate of calcium events in both brain regions, as selectivity depends on neuronal activity levels. In both V1 and the dentate gyrus, the overall level of activity was markedly increased during locomotion (Fig. 2a), as previously reported in other studies (Niell and Stryker, 2010; Pilz et al., 2016). This increase occurred in a step-like manner, i.e. it depended on the animal being in motion or standing still, but did not appear to further correlate with speed once the animal was moving (Fig. 2c). While activity levels were similar between V1 and DG during rest, V1 was significantly more active during running periods (Fig. S2a,b). Notably, there was no difference in event rate between active and passive conditions during running in both V1 and DG (Fig. 2b). The decoupling of running speed from VR motion during passive exposure allowed us to test whether this increase was arising from motor activity, or if it reflected the difference between static and moving visual inputs. In V1 but not in DG, the event rate was increased when the projector displayed motion compared to periods of static images (S2c,d). However, the animals also ran more when the visual flow was moving (S2e,f), preventing us from further refining this analysis. Overall, these results indicate that neuronal activity rates in both the visual cortex and the dentate gyrus depend on locomotion but not on behavioural engagement.

**Figure 2:**
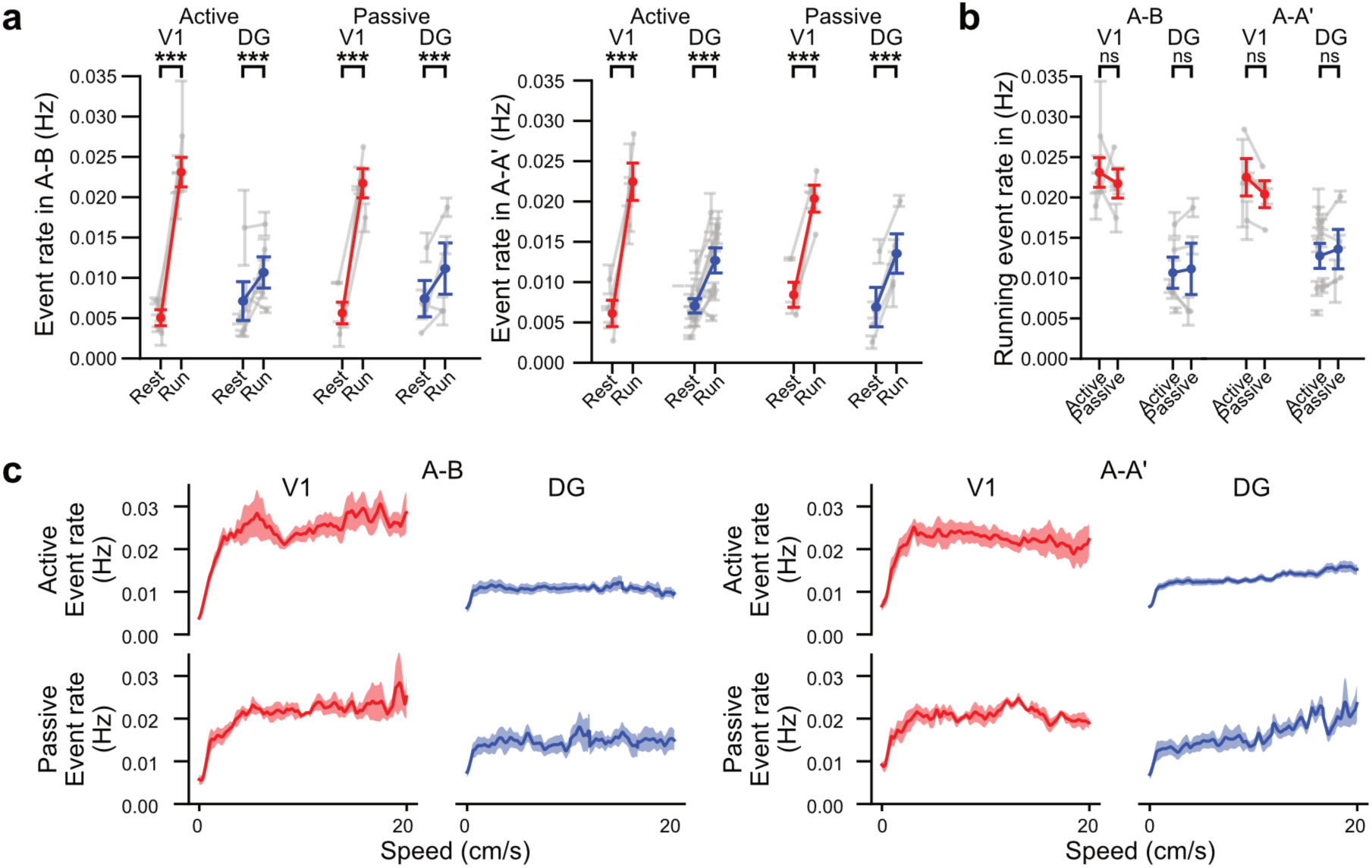
Two-photon calcium imaging reveals that activity levels are driven by locomotion but not by behavioral context in V1 and DG. **(a)** Comparison of the event rate between resting and running for the A-B environment pair (left) and the A-A’ environment pair (right). **(b)** Comparison of event rate during running between active and passive conditions in A-B (left) and A-A’ (right). Same values as in **(a)**. **(c)** Event rate as a function of running speed for the A-B pair of environments (left) and the A-A’ pair (right). In **(a)** and **(b)**, error bars indicate the standard error of the mean. Grey lines correspond to multiple sessions from a single animal, while coloured lines show the average across animals. In **(c)**, solid lines show the mean while shaded areas correspond to the standard error of the mean.

To assess neuronal discrimination of sensory inputs in the visual cortex and in the dentate gyrus in a time-resolved manner at the population level, we employed a binary decoder to predict the explored environments from calcium signals. In brief, we trained a CEBRA model to project data points, i.e. vectors representing the level of activity across each neuron, into a three-dimensional space while maximising the distance between samples from different environments (Fig. 3a) (Schneider et al., 2023). Subsequently, we separated the testing data into two groups using k-means clustering (Fig. 3b). Employing a cross-validation strategy, we constructed an ensemble decoder and generated a prediction signal that covered all time bins throughout the session, representing the consensus classifications derived from multiple data partitions (Fig. 3c). The chance level of decoding was calculated by randomly splitting lap assignments, which revealed that the decoder systematically performed above the chance level in the visual cortex. However, in the dentate gyrus, decoding was only successful when the animals were actively navigating (Fig. 3d, Fig. S3a). We could also confirm these findings based on the selectivity of individual neurons (Fig. S4), suggesting that most of the information about the explored environment is already encoded at the single-neuron level.

**Figure 3:**
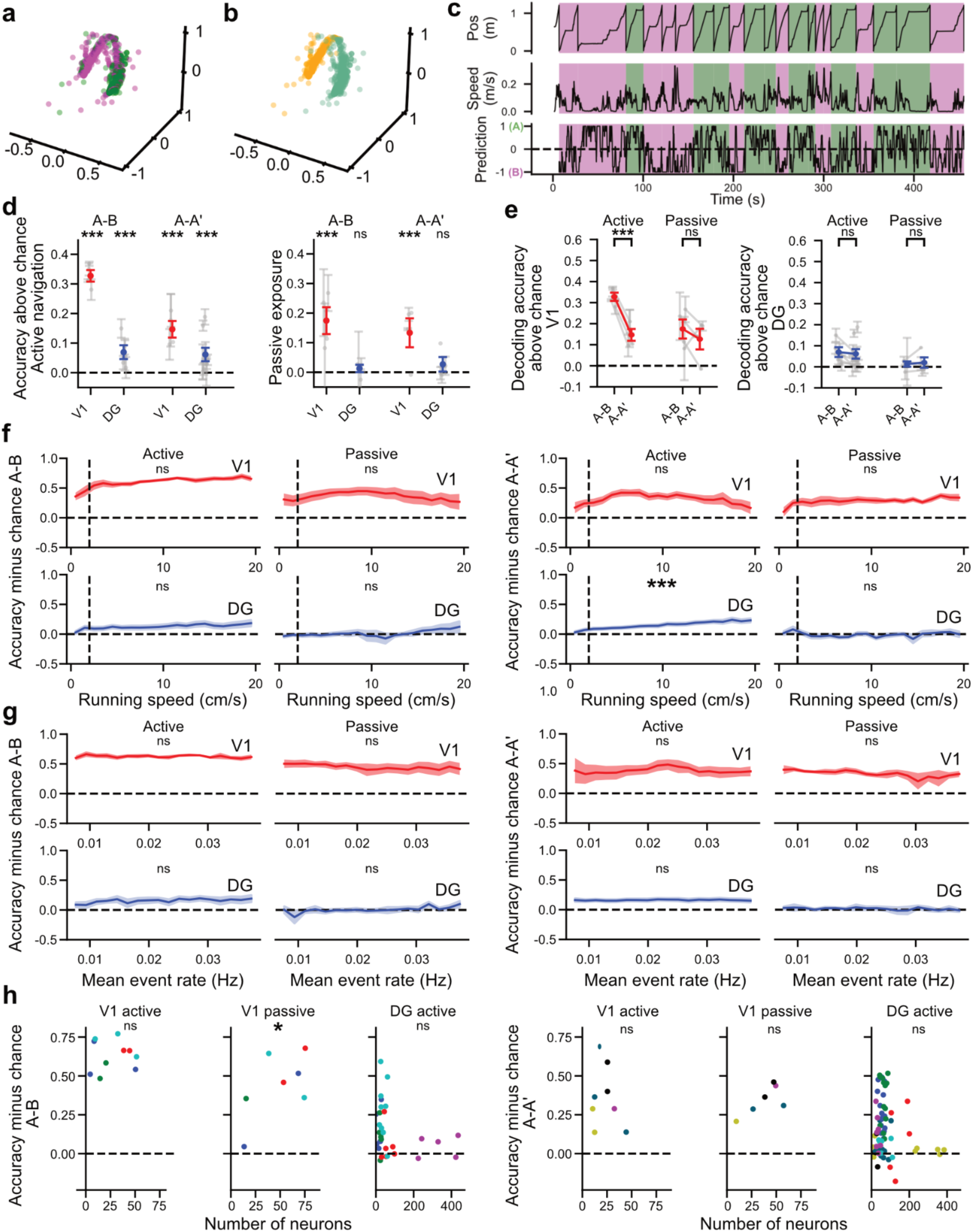
Visited environments can be decoded from neuronal activity only during active navigation in the DG, but under both active and passive conditions in V1. **(a)** Three-dimensional projection of neuronal activity using the CEBRA algorithm. Each point corresponds to a time bin within the recording. Time bins are coloured based on the visited environment, i.e. the ground truth (green: A, magenta: B) **(b)** Same as (a), with time bins coloured based on the cluster assignment from a k-means clustering, i.e. the prediction. **(c)** Example ensemble decoder obtained using a cross-validation strategy. +1 indicates a perfect consensus across all predictions for environment A, whereas -1 corresponds to a perfect consensus for environment B. **(d)** Decoding accuracy with chance level subtracted for active (left) and passive (right) conditions. **(e)** Comparison of decoding accuracy between both pairs of environments in V1 (left) and DG (right). **(f)** Decoding prediction as a function of running speed for the environment pair A-B (left) and A-A’ (right). Linear regression is only computed above a threshold set at 2 cm/s (dashed line) as activity is too sparse at low speed (Fig. 2D). **(g)** Decoding accuracy as a function of the event rate for environment pair A-B (left) and A-A’ (right). **(h)** Decoding accuracy as a function of the the number of segmented neurons for environment pair A-B (left) and A-A’ (right). In **(d,e)**, error bars indicate the standard error of the mean. Grey lines correspond to multiple sessions from a single animal, while coloured lines show the average across animals. In **(f,g)**, solid lines show the mean while shaded areas correspond to the standard error of the mean. In **(h)**, each dot corresponds to a single session, colours represent animals.

The dentate gyrus has been suggested to perform pattern separation (Allegra et al., 2020) by orthogonalising input patterns. To assess whether our analysis could confirm this notion, we compared the decoder performance between both pairs of environments during the same behavioural paradigm (Fig. 3e). While decoding accuracy was higher in the visual cortex for the more distinct pair of environments A-B, no significant difference in accuracy was found in the dentate gyrus between the two pairs of environments. Thus, our results are consistent with the view that orthogonalisation in the dentate gyrus occurs even when the differences between the environments are small. Interestingly, the increased discrimination in the visual cortex appears to be specific to active navigation, as there was no difference upon passive exposure.

Hippocampal place cells tend to cluster around landmarks and reward zones, giving rise to an “overrepresentation” of these spatial locations (Gauthier and Tank, 2018; Hollup et al., 2001; Sato et al., 2020) with increased spatial selectivity and stability (Bourboulou et al., 2019). To test whether neuronal discrimination is enhanced in particular locations along the track, we visualised the decoding accuracy as a function of the position on the track (Fig. S3c,d). This analysis revealed that neuronal discrimination was homogeneously distributed along the track. In particular, the decoder did not perform better within reward zones (Fig. S3e,f). This finding might partly be explained by the insight that the mechanisms underlying representations of distinct environments are more closely related to remapping processes (Fyhn et al., 2007) than to positional decoding, and thus the location does not influence the decoder performance unless the environment changes.

Since the forced behavioural disengagement induced by passive exposure had a marked effect on the level of neuronal discrimination, we sought to determine whether we could relate behavioural metrics, which may reflect more subtle fluctuations in behavioural engagement, to decoding accuracy while animals are actively performing the task. First, we investigated a potential correlation between decoding accuracy and locomotion. To this end, the predictions from the decoder were divided into bins of animal speed that were linearly spaced between 0 and 20 cm/s. A visual inspection of the data suggested a potential correlation between decoding accuracy and running speed in the dentate gyrus. As had been observed with the event rate, at very low running speeds, the decoder accuracy drastically dropped (Fig. 3f). Consequently, a cutoff was set and only bins with a running speed higher than 2 cm/s were considered for statistical analysis. To formally test this hypothesis, a linear mixed model was employed to relate decoding prediction to speed, with sessions treated as a statistical unit. In V1, we found no correlation between running speed and decoder accuracy. In contrast, in the DG, the decoder performance was positively correlated with running speed for the A-A’ pair of environments, but not for the more distinct A-B pair. Moreover, we found no correlation between the decoder accuracy and the speed of the visual flow during passive exposure (Fig. S3b). This result suggests that in the hippocampus, decoding accuracy may be more closely related to the behavioral state of the animal, such as the level of alertness or running, than in the visual cortex. We also assessed the relationship between the behavioural and decoder performances using the ratio of anticipatory licks, but could not identify any significant difference (Fig. S3g,h). Importantly, the differences between the DG and V1 were not caused by the sparsity of the DG or the number of neurons within the field of view, as the decoder performance overall did not correlate with the event rate (Fig. 3g) or the number of recorded neurons (Fig. 3h).

In summary, our findings demonstrate that the visual cortex consistently produces faithful representations of the environment, independent of active engagement in behavioral tasks. Conversely, neuronal representations in the hippocampal dentate gyrus depend critically on the behavioral relevance of visual inputs, discriminating between virtual environments only during active participation in the task. These results reveal a fundamental distinction in sensory processing between primary sensory cortices and the hippocampus, reflecting their distinct roles in encoding sensory and spatial information.

Our direct comparison of sensory processing in the visual cortex and the hippocampus under identical experimental conditions reveal key differences in how visual and spatial information is processed in these two brain regions. The visual cortex reliably distinguishes visually richer environments more effectively than sparse ones, irrespective of the behavioral context, revealing its role in providing relatively neutral sensory representations. In contrast, the dentate gyrus does not differentiate between visually rich and sparse environments based on visual complexity alone but instead generates orthogonal representations driven by task relevance. This aligns with previous evidence that the dentate gyrus provides decorrelated sensory inputs to downstream hippocampal circuits for cognitive mapping (Allegra et al., 2020; Gómez-Ocádiz et al., 2022).

Importantly, our results do not question a large number of studies showing that visual cortex is subject to top-down influences (Gilbert and Li, 2013; Kanamori and Mrsic-Flogel, 2022; Niell and Stryker, 2010; Pakan et al., 2016; Zhong et al., 2025) . Rather, we extend this concept by showing that in comparison to a higher-order brain region, V1 provides a base representation of the visual environment that is independent of the behavioral context. However, already at this early stage of sensory encoding, our results also confirm that in addition to this robust base representation, behavioural context modulates neuronal responses (e.g. Fig. 2a).

The behavioral dependence of hippocampal representations is also supported by the positive correlation between running speed and neuronal discrimination performance in complex tasks such as distinguishing between the two similar environments A-A’. This observation confirms previous findings that spatial signals and neuronal activity in the hippocampus are strongly influenced by behavioral states, active navigation and rewards (Allegra et al., 2020; G. Chen et al., 2013; Pettit et al., 2022; Sosa et al., 2025; Terrazas et al., 2005). Although visual cortical neurons also exhibit spatial modulation during active navigation (Flossmann and Rochefort, 2021; Haggerty and Ji, 2015; Pakan et al., 2018; Saleem et al., 2018), our data show that spatial representations are less dependent on behavior than in the hippocampus and remain robust even during passive exposure.

Additionally, our analysis demonstrates that spatial coding and neuronal discrimination can be dissociated. For instance, passive exposure led to reduced spatial information in both the visual cortex (consistent with (Diamanti et al., 2021)) and the hippocampus, but entirely abolished neuronal discrimination only in the hippocampus. This indicates that spatial representation alone does not necessarily equate to environmental discrimination, further reinforcing the specialized roles of these regions.

Our findings demonstrate that task relevance is encoded already at the earliest hippocampal processing stage, the dentate gyrus. While we have previously shown that representations in the dentate gyrus are not predictive of task performance (Allegra et al., 2020), here we reveal that they depend on active participation in a task, effectively functioning as a gate that selects sensory information based on behavioral engagement. By contrast, representations in CA1 also correlate with task performance, thereby encoding task outcome (Allegra et al., 2020). These findings suggest a hierarchical model of sensory processing, where sensory cortices act as continuous input providers, the entry stage of the hippocampus acts as an on-off-switch for task relevance, and the output stage encodes task outcome.

## Methods

### Animals

All procedures were carried out in accordance with European and French guidelines on the ethical use of animals for experimentation (EU Directive 2010/63/EU) and were approved by the Ethics Committee CETEA of the Institut Pasteur (protocol numbers 160066 and 220037). Adult male C57BL/6J wild-type mice (Janvier Labs) were employed for all experiments. The mice were five weeks old at the time of arrival and underwent a one-week acclimation period in the animal facility prior to the commencement of surgical procedures at six weeks of age. Mice were housed in a room kept at 21°C in groups of two to four in polycarbonate individually ventilated cages, which were enriched with running wheels. The mice were kept on a 12-hour inverted light/dark cycle and were provided with ad libitum access to food and water. Depending on the type of implants, either a single or two surgeries, separated by two weeks, were performed on the animals (see surgery section). The training protocol typically commenced between two to six weeks after the last surgeries, therefore on 8- to 14-week-old animals. All experiments were conducted during the dark phase of the light cycle. During the experimental protocol, animals were typically housed in pairs, although exceptionally they could be single-housed to end fighting, or because of the death of their littermate.

Part of the data from the dentate gyrus were obtained from animals that were also used for a previous publication; see in particular Fig. S4I,J (Allegra et al., 2020).

### Virtual reality navigation

#### Setup

We used a one-dimensional VR system with a spherical screen. Animals were connected via their headposts to a head-fixation system positioned right above a 20 centimetres diameter polystyrene cylindrical treadmill covered by gaffer tape. The cylinder could rotate bidirectionally along a single axis with bearings guaranteeing a sufficiently low friction so the strength exerted by a running adult mouse would easily put it in motion. We positioned the sensor from a computer gaming mouse (G203 from Logitech) at close proximity (a few millimetres) of the cylinder to measure its rotation along a one-dimensional axis with an acquisition frequency of 1 KHz.

We then used the Blender Game Engine (http://www.blender.org) together with the Blender Python API to convert the rotation speed into a new position within a virtual linear corridor at a 100 Hz sampling rate. During each individual epoch of 10 ms, the position in the VR system was incremented by the distance covered. A new frame was rendered at the novel coordinates and displayed back to the animal for another 10 ms while treadmill rotation kept being acquired, hereby forming a closed loop system. To display the visual flow arising from these frames, we used a projector (Casio XJ-A256) pointed towards a quarter sphere mirror of 45 cm located under the mouse. The mirror reflected back the video onto a 120 cm diameter spherical dome screen wrapping a 240°angle around the animal which covers almost its entire field of view.

For training purposes, a reward delivery system was implemented. Tubing was positioned within the licking distance of mice using Thorlabs components. A pump controlled by the software would deliver a drop of sugar water to drop (10 μl, 8 mg/mL sucrose) as a reward while a piezo sensor would record single licks from the animal.

#### Environments

Three virtual linear corridors were employed for the discrimination tasks. The gain was calibrated so that the length of each corridor corresponded to a physical distance of 1.28 metres run on the cylindrical treadmill. No specific cue was displayed to indicate the location of the reward zones; however, numerous proximal cues were distributed along the length of the track, which may be employed by animals to guide navigation. In order to ensure that animals were required to discriminate between environments, two locations were employed for the reward zones. In A, the reward zone was located between 0.85 m and 1.20 m. In both A’ and B, that zone spanned from 0.35 m to 0.71 m. The delivery of the reward was contingent upon the animal spending at least three seconds within the reward zone.

Animals were initially trained exclusively on environment A, which can be considered to be the familiar environment given that they have been largely more exposed to this one than to the two others. Conversely, the environments A’ and B are considered novel. The environment A’ strongly resembles A, with the exact same proximal cues at the same positions, and only differs by the grating on the wall, which follows a vertical angle in A and an oblique orientation in A’. Environment B was markedly distinct from A/A’ in terms of visual richness, with entirely distinct proximal cues that were in a greater number than in previous environments.

#### Active navigation

Approximately one month following the surgical procedures, the animals were placed under a two-photon microscope to check how well the calcium indicator was expressed. Animals displaying promising signals were selected for behavioural training prior to imaging. In brief, they were placed under water restrictions and handled for approximately ten minutes daily for three consecutive days. Subsequently, the animals were head-fixed twice on the behavioural setup for 10 to 20 minutes, without any environmental stimuli being presented, in order to habituate them to the protocol and to the restriction of their head movements. They were then trained to navigate the single corridor A for five sessions of 30 minutes. Laps could be rewarded or not, depending on the amount of time spent within the reward zone. In the event that an animal failed to trigger the delivery of a reward, no additional punishment was administered. The only consequence was the time lost by the animal to complete the lap, i.e. no timeout, air puff, sound or light flash was employed. When reaching the end of a lap, animals would cross a black vertical wall and be immediately teleported back to the beginning of the track. Recording sessions lasted approximately 500 seconds, during which two environments were alternated at random, with a 50% probability of each environment being displayed at the beginning of every lap. Each session could be either a ‘similar session’, when the environments A and A’ were alternated or a ‘distinct session’, when A and B were presented in alternation. Environments A’ and B were never alternated.

In most cases, two sessions were recorded per animal within a single day, although on few occasions up to four recording sessions were conducted. The animals were placed on the behavioural setup and trained on the task on a daily basis. However, to reduce the risk of bleaching, they were usually recorded only every other day. The recording period typically lasted for approximately three weeks. In order to maintain motivation, animals were subjected to water restriction from the beginning of the training protocol. During the week, they had only access to water via the reward deliveries and a supplementation of HydroGel (from ClearH2O). Their body weight was not allowed to fall below 80% of their pre-restriction value. The quantity of HydroGel provided to the animals was dependent on their weight. They could receive between 0.3 g and 1.2 g daily, with a typical amount of 0.5 g. During weekends, 5 mL of water was provided to each animal. Results from 20 mice are included in this study (V1: n = 4, DG: n = 9, CA1: n = 7).

#### Passive exposure

In order to test our hypothesis, we required a condition in which animals were exposed to similar sensory inputs that would be devoid of behavioural relevance. For this purpose, we decided to use passive exposure. Rather than inferring the position within the VR system from the rotation speed of the wheel, we decoupled these two variables. The position was obtained by re-running an old session, which determined both the displayed visual flow and the delivery of rewards. The animal was not restrained in its locomotion, yet it was unable to control any aspect of the VR system. The same replayed session was used for all passive exposure recordings. Importantly, only the locomotion and reward delivery were extracted from the replayed session. The environment displayed was randomly selected between either A-A’ or A-B, depending on the condition. This implies that, although rewards were delivered within one of the two reward zone locations, this location was not necessarily the one associated with this environment. To say it differently, a lap initially performed within environment A may be displayed during the replay as A’ or B, although it will show the animal stopping at the reward zone of A; and vice versa. The design of the task was thus chosen to ensure that the visual flow was behaviourally irrelevant, as otherwise it would allow the animal to predict the timing of reward delivery.

### Surgical procedures

#### General procedures

Surgical procedures were performed on a stereotaxic frame (Kopf Instruments). Prior to the beginning of the surgical procedure, a combination of buprenorphine (0.05 mg/kg, Vetergesic) and meloxicam (10 mg/kg, Metacam) was administered intraperitoneally, 30 minutes before to the surgical incision. The opening site was meticulously shaved using a hair clipper and cleansed with a solution of povidoneiodine (Betadine). Local anaesthesia was ensured by infiltrating lidocaine for a period of 30 seconds prior to the initial incision. For the induction of anaesthesia, mice were exposed to isoflurane at a concentration of 3-5%. During the surgical procedure, the animals were maintained anaesthetised with a 1-2% isoflurane mixture, while their body temperature was kept at 36°C using a heating blanket and their eyes were protected with artificial tear ointment. Identification of the animals was achieved by marking their ears during the surgical procedure. Postoperative care involved the oral administration of meloxicam (5 mg/kg) mixed with surgical recovery DietGel (from ClearH2O) for two consecutive days.

Following the incision, the exposed skull was cleared of connective tissues. A craniotomy was performed above the region of interest. For hippocampal implants, the dorsal region in the right hemisphere was targeted, with the craniotomy centred 2 mm posterior and 1.5 mm lateral from the Bregma reference point. In the visual cortex, the craniotomy was centred 3 mm laterally from the Lambda reference point, in the right hemisphere. An adeno-associated viral construct (AAV) carrying the GCaMP6f indicator (T.-W. Chen et al., 2013) (AAV1.Syn.GCaMP6f.WPRE.SV4, titer 3.4x10^12^ TU/mL; Addgene) was injected at the centre of the craniotomy via a glass micropipette (Wiretrol, 5-000-1010 Drummond). For hippocampal recordings, 500 nL of virus were injected over an 8-minute period at a depth of 1.7 mm from the dural surface for dentate gyrus implants and 1.2 mm for CA1 animals. In the visual cortex, 900 nL of virus was injected as three thirds at 900, 500 and 300 µm depth, over a 15-minute period.

Subsequently, three distinct implant types could be employed, each requiring a different surgical technique. While the visual cortex windows were implanted simultaneously with the viral injection, the craniotomies above the hippocampus were closed using surgical glue and the optical implantation procedure was performed one to two weeks later using the same anaesthesia protocol as described above. Regardless of the implant type, a stainless-steel headpost for head fixation (Luigs & Neumann) was positioned at the end of the final surgery and maintained in position using dental cement (Super-bond C&B, Sun Medical). Protective silicone elastomer was positioned on top of the implant for protection (900-2822, Henry Schein), removed and repositioned after each recording session.

#### Cannulas

Cannula implants were preferentially employed for CA1 recordings. The cannulas were manually assembled using UV glue (Norland optical adhesive) to attach a circular cover glass (1.6 mm diameter, 0.16 mm thickness, produced by Laser Micromachining) to a cylindrical stainless-steel tube (2 mm height, 1.65 mm outer and 1.39 mm inner diameter; Coopers Needleworks). A new craniotomy of 1.6 mm diameter was drilled, centred on the previous one. The overlying cortex, which is composed primarily of somatosensory areas and posterior parietal association cortices, was aspirated using a 27-gauge needle connected to a pumping system. During aspiration, brain tissue was constantly rinsed using an aCSF (artificial cerebrospinal fluid) solution, while a dental sponge was employed to control bleeding. For CA1 recordings, aspiration was terminated when the external capsule became visible, thus preserving the integrity of the hippocampal formation. For DG recordings, it was necessary to remove the CA1 area located directly above the DG, in addition to the cortex. Following the cessation of aspiration, the cannula was slowly pushed into the brain, slightly deeper than the injection depth, that is about 1.4 mm deep for CA1 and 2 mm deep for DG implants. This was required as the applied pressure also displaces the surrounding tissue. Protruding parts of the cannula were then secured to the skull with opaque dental cement (Super-bond C&B, Sun Medical) (Allegra et al., 2020).

#### GRIN lens

As an alternative strategy to reduce tissue lesions, we implanted gradient-index (GRIN) lenses (1 mm diameter, 3.4 mm height, NA = 0.5, G2P10 from Thorlabs), which were preferentially used for DG recordings as these surgeries are more invasive than for CA1. A new craniotomy of 1 mm diameter was drilled, centred on the previous one. In lieu of cortical aspiration, a bevelled stainless-steel cylinder (1 mm diameter) was lowered to the targeted region using the stereotaxic frame. It was then removed, before slowly inserting the GRIN lens into position, at the same coordinates as the aforementioned cannulas. GRIN lenses were also maintained in position by applying dental cement to the protruding parts.

#### Cortical windows

In order to image from the visual cortex, a circular cover glass (3 mm diameter, 0.16 mm thickness) was implanted at skull level. A single craniotomy (3 mm diameter) was drilled at the beginning of the procedure and used both for viral injection and implant positioning. As this surgery does not entail the removal of any cortical tissue, the craniotomy was executed with great care to avoid damaging the dura or causing bleeding. The glass window was then positioned on the exposed cortex and pressed tightly onto the brain using the stereotaxic arm. Finally, opaque dental cement was applied to the edges of the glass window and to the skull in order to seal them together

### Tissue histology

Once the experiments had been completed, the animals were euthanised and their brains were recovered for post-hoc histological confirmation of the recorded brain area. First, they were deeply anaesthetised with an overdose of ketamine/xylazine administered intraperitoneally. Subsequently, following the cessation of reflex responses, the animals were perfused transcardially with phosphate-buffered saline (PBS, 1x) to wash out the blood circuit, followed by 4% paraformaldehyde (PFA) solution for tissue fixation. The brains were then extracted and kept in 4% PFA solution overnight, before being cut into 60-70 µm thick coronal slices the next day. The slices were stained with DAPI in order to use the nuclei as general landmarks for neuroanatomy. Sections underneath the optical implant were collected and mounted for fluorescence microscopy, where we could confirm both which population of cells expressed the GCAMP indicator and where the implant was exactly positioned. Fluorescence images were acquired with a spinning disc confocal microscope (Opterra, Bruker).

### 2-photon calcium imaging

Prior to any recording sessions, the protective silicone was removed and the implants were cleaned using distilled water and an air spray. The recordings were performed using a resonant-galvanometer high-speed laser scanning two-photon microscope (Ultima V from Bruker) in conjunction with a 16x 0.8 NA water immersion objective from Nikon. This permitted the acquisition of time series at a frame rate of 30 Hz, with a field of view of 512x512 pixels, each pixel being approximately 1 µm². We employed a femtosecond-pulsed excitation laser (Chameleon Ultra II, Coherent) tuned to 920 nm for imaging the GCaMP6f calcium indicators. To prevent the light emitted by the VR projector from interfering with the fluorescence measurements, a black foam rubber ring was positioned between the animal’s implant and the objective. Additionally, a green light-blocking filter (FES0450, Thorlabs) was placed in front of the projector light output.

### Extracting neuronal activity

The built-in algorithm from the Suite2p software was employed for motion correction and cell segmentation (Pachitariu et al., 2017). In brief, motion correction was performed using a nonrigid registration technique. An initial reference image was generated from the frames sharing the highest correlation between each other. Then, individual frames were shifted to maximise their alignment to this reference frame. Instead of cross correlation, this algorithm relies on phase-correlation, which normalises the Fourier spectra of both images prior to multiplying them. As a non-rigid registration was applied, each image was further subdivided into small blocks, which were individually shifted to maximise correlation to the reference image.

Cell segmentation was achieved through singular value decomposition, a matrix factorisation technique that shares similarities with principal component analysis (PCA). In order to identify spatial components, the movie was binned and the mean signal across bins was subtracted. Subsequently, the movie was smoothed across the X/Y axes using a Gaussian filter and each pixel was normalised by its own variance. Finally, the covariance matrix of the movie was computed and factorised using singular value decomposition. The top spatial components were retained for further computations. A correlation map was generated by smoothing the spatial components across the X/Y axes and dividing the smoothed components by the unsmoothed ones. Subsequently, peaks were identified in the correlation map and regions of interest (ROIs) were iteratively extended around them. The signal was extracted by calculating the mean fluorescence within each ROI mask. Finally, the neuropil in the vicinity of each cell was computed using raised cosines as a spatial basis function. The segmented cells were manually curated based on both their anatomy and fluorescence traces. In the dentate gyrus, ROIs with large isolated cell bodies were excluded in order to prevent the inclusion of putative interneurons and mossy cells. Instead, we specifically selected ROIs which had small densely packed cell bodies. Cells exhibiting clear calcium transients followed by exponential decay were selected.

Afterwards, normalised fluorescence traces were extracted by subtracting the neuropil signal, scaled by a factor of 0.7, from the ROI mean fluorescence. Given the clear and distinct calcium transients exhibited by the cells, followed by a return to stable baseline, we elected to employ an event detection method in lieu of a deconvolution approach. This was because event detection requires fewer assumptions and little information would be lost. In particular, we were concerned that a deconvolution approach that we would not be able to calibrate carefully with ground truth data may introduce a bias when comparing between regions, given that the principal cell populations in V1, DG and CA1 have substantially different characteristics. Calcium events were defined as continuous time periods, each lasting at least 300 ms, during which the normalised fluorescence deviated from the mean value by more than 2.5 standard deviations (Allegra et al., 2020; Dombeck et al., 2010). Each event was then characterised by a unique timestamp value, which was localised at the midpoint of the rising phase of the transient.

While a subset of neurons with stable spatial modulation of firing (“place cells”) could be found in both V1 and in the DG in our data set (Figure S5), all analyses were performed on all cells (place cells and non-place cells) to avoid discarding cells that contribute to spatial coding without apparent spatial modulation of their activity (Stefanini et al., 2020).

### Selectivity

The most basic form of neuronal discrimination is the preferential firing of neurons for one of the two environments. To formally quantify this bias, we employed selectivity. This metric is a signed value obtained for each neuron individually by computing the signed difference between the mean event rate in environment A and the mean event rate in environment A’/B. The value is normalised by dividing by the sum of the mean event rate in both environments, thus resulting in a value bounded between -1 and +1. A negative value indicates a preferential firing in the novel environments (A’/B), while a positive value corresponds to a higher activity within the environment A. An absolute value of one implies that the neuron exclusively got activated in the corresponding environment, while a neuron with a selectivity of 0 has perfectly identical activity in both corridors. The formula employs event rate rather than the number of events in order to account for the time spent within each environment. Due to the marked effect that locomotion had on the activity levels, selectivity was computed based only on time periods with locomotion.

At the population level, selectivity was defined as the mean value of the absolute selectivity of each neuron from a given recording session. Absolute selectivity does not indicate towards which environment the neuronal activity is biased; rather, it reflects the degree of preferential firing exhibited by the population. Importantly, the absolute selectivity calculated from a pure noise signal is largely above 0, given that recordings are constrained by a finite duration and number of events. For instance, a neuron that fires twice may do so once in environment A and once in environment B, but it may also fire twice in either environment. In a recording in which the same amount of time is spent in both environments, a neuron with two calcium events has an expected value for selectivity of 0.5. To address this issue, the chance level for each session was calculated by computing selectivity on surrogate data. To maintain the task structure of the recordings, the surrogate data was obtained by randomly shuffling the labels of laps as A or A’/B, while maintaining the number of laps in each environment. This operation was repeated 100 times, and the chance selectivity was obtained by averaging the outcome of these 100 iterations.

### Decoder

To quantify more rigorously how informative the population activity was about the environment, we employed a binary decoder to predict the explored environments based on the activity of all neurons within small temporal bins of 33 ms. In brief, the binary activations were initially smoothed by applying a one-dimensional Gaussian filter (500 ms standard deviation) to the vector of binary activations. Subsequently, in order to rescale the signal, a multiplication by the sampling rate was performed. In other words, each binary activation was replaced by a Gaussian kernel with a standard deviation of 500 ms and a peak scaled at approximately 0.78.

Laps were divided into two halves, separately for each environment, for the purposes of training and testing the model. A cross-validation strategy was employed, with 20 random partitions of the data being performed to ensure that each lap was used for both training and testing. In order to guarantee a minimum level of activity and thus enable the model to make meaningful predictions, a threshold was set, requiring that the sum of the smoothed activity across neurons exceed 0.3 in a time bin. During periods when a single neuron was active across the entire population, this threshold excluded time bins that were further than 1.5 standard deviations from the event, i.e. ± 750 ms.

The clustering approach was based on the recently published framework CEBRA (Schneider et al., 2023). It is designed to project neuronal activity into a low-dimensional manifold based on behavioural variables. The model was trained to perform three-dimensional projections in the CEBRA-Behaviour mode, with the explored environment inputted as an auxiliary variable. Subsequently, the testing data was projected using the trained model. If the environments were represented in the data, two clusters should emerge, corresponding to the explored corridors. k-means clustering with two clusters was therefore performed in the low-dimensional space. Importantly, in contrast to the typical approach, which involves first applying an unsupervised dimensionality reduction algorithm (such as principal component analysis, independent component analysis, or t-SNE) and then performing supervised clustering in the low-dimensional space, the method presented here employs a reversed sequence. The latent embedding is supervised and trained with the objective of maximising the separation between the two clusters, whereas the clustering algorithm is unsupervised. Consequently, the cluster labels are abstract and therefore required to be reassigned to the environments. To achieve this, both assignments were tested and the combination that maximised accuracy was selected. As a result of this best-matching strategy, although the classification is binary, the chance level is slightly above 50%.

Subsequently, the 20 partitions of the data were combined by averaging in order to obtain a prediction signal across the entire recording. This signal reflects the confidence of the decoder, as it is based on the consensus from all the decoders that made a prediction for this particular time bin (on average, 10 out of the 20 decoders, although this fluctuates based on the random partitioning). A value of 1 indicates that all decoders predicted environment A, while a value of -1 reflects a perfect agreement for the novel environment (A’ or B, depending on the condition). As a minimum activity threshold was set, no predictions were made for time bins that did not meet the requisite criteria.

Surrogate data was generated to formally assess the chance level by random shuffling lap assignment, in a similar manner to selectivity. The number of laps within each specific environment was maintained. The same computing pipeline as with the unmodified experimental data was enforced. Laps labels (as A or A’/B) were shuffled once, then 20 random partitions of the data were generated and combined into a single ‘chance’ decoder. This analysis was repeated 10 times, with a different random labelling of laps for each iteration. In total, 220 decoders based on data partitions were built for each recording session, and 11 decoders were generated by averaging across partitions (one from the original data and 10 from the shuffled data).

Evaluation of the model performance was performed on the ensemble decoder obtained after the merging of partition outcomes. We used the confidence-weighted accuracy, simply referred to as accuracy in the text and figures. In brief, instead of computing the accuracy in a binary manner as correct/incorrect classification, each prediction was weighted by its corresponding confidence (i.e. consensus across partitions).

### Behavioural metrics

For the behavioral analyses, more animals were included than in the neuronal activity analysis, as calcium activity from certain animals was unusable due to various factors, including poor signal quality and targeting of incorrect brain regions. Each session was divided into running and resting periods based on the speed of the animal. The transitions between the running and resting states were identified through the implementation of a threshold strategy. A cutoff value of 0.5 cm/s was defined, and whenever the speed exceeded or dropped below this threshold for a period of at least one second, this resulted in the commencement or cessation of a running or resting period. In order to compute the motion of the visual flow, the derivative from the position in the VR system was calculated and low-pass filtered in a manner similar to that applied to the physical speed sensor. In some sessions, the sensor malfunctioned, resulting in unusually elevated lick rates. Based on the distribution of mean lick rate across all recordings, a threshold was set and sessions with a mean lick rate greater than 5 Hz were excluded from analyses involving licks (anticipatory licks, hit rate). This represented 23% of sessions. Anticipatory licks were defined as licks occurring within a two-second window prior to the delivery of a reward. Consumption licks were defined as licks occurring within a five-second window following the delivery of a reward. A hit was defined as a reward being delivered with at least three anticipatory licks displayed.

## Acknowledgements

We thank Nathalie Rochefort and Tom Flossmann for helpful discussions and comments on the manuscript. We thank Lucile Sontag and Claire Lecestre for technical assistance. This work was supported by grants from the ERC (StG 678790 NEWRON to C.S.-H. and MSCA 800027 FindMEMO to M.A.), the Agence Nationale de la Recherche (ANR-22-CE37-0008 VIPLANAV to C.S.-H.), the Fondation pour la Recherche Médicale (FDT202304016565 to C.O.), the Fondation des Treilles (to C.O.) and the INCEPTION programs D2I and S2I (Investissement d’Avenir grant ANR-16-CONV-0005 to C.O.). The Investissement d’Avenir grant ANR16-CONV-0005 funded computing resources used for the decoding analysis.

## Author contributions

M.A. and C.S.-H. conceived the project. C.O., M.A. and C.S.-H. designed experiments. C.O. and M.A. performed the experiments. C.O. analysed data. C.S.-H. supervised the project. C.O., M.A. and C.S.-H. wrote the manuscript.

## Statistics (main figures)

All statistical values are compiled in a dedicated table at the end of the figure section.

LMM: Linear Mixed Model; TT: paired t-test; MW: Mann–Whitney U test; WIL: Wilcoxon signed-rank test; BENF: Benferroni correction for multiple comparisons. When providing N values for linear mixed models, the first row corresponds to the number of groups and the second row to the number of observations.

IAV production of the monoclonal MOCK celllines CS9 andC113.

| Panel | Field 1 | Value | N | Field 2 | Value | N | Test | p-val |
| --- | --- | --- | --- | --- | --- | --- | --- | --- |
| **Figure 1** |  |  |  |  |  |  |  |  |
| 1g left | Run time A-B active | 68.1 ± 3.4 | 16 animals<br>145 sessions | Run time A-B passive | 75.3 ± 5.4 | 11 animals<br>28 sessions | LMM | 0.004 |
| 1g right | Running speed A-B active | 10.3 ± 0.5 | 16 animals<br>145 sessions | Running speed A-B passive | 13.1 ± 2.3 | 11 animals<br>28 sessions | LMM | 0.002 |
| 1i | Anticipatory licks A-B active | 9.1 ± 1.4 | 16 animals<br>141 sessions | Anticipatory licks A-B passive | 6.4 ± 0.7 | 11 animals<br>29 sessions | LMM | 0.030 |
| **Figure 2** |  |  |  |  |  |  |  |  |
| 2a left A-B | Active V1 Run | 0.005 ± 0.001 | 4 animals<br>10 sessions | Active V1 Rest | 0.023 ± 0.002 | 4 animals<br>10 sessions | LMM | 3e-17 |
| 2a left A-B | Active DG Run | 0.007 ± 0.002 | 5 animals<br>30 sessions | Active DG Rest | 0.011 ± 0.002 | 5 animals<br>30 sessions | LMM | 0.001 |
| 2a left A-B | Passive V1 Run | 0.006 ± 0.001 | 4 animals<br>7 sessions | Passive V1 Rest | 0.022 ± 0.002 | 4 animals<br>7 sessions | LMM | 2e-23 |
| 2a left A-B | Passive DG Run | 0.007 ± 0.002 | 4 animals<br>14 sessions | Passive DG Rest | 0.011 ± 0.003 | 4 animals<br>14 sessions | LMM | 6e-6 |
| 2a right A-A' | Active V1 Run | 0.006 ± 0.002 | 4 animals<br>8 sessions | Active V1 Rest | 0.022 ± 0.002 | 4 animals<br>8 sessions | LMM | 9e-9 |
| 2a right A-A' | Active DG Run | 0.007 ± 0.001 | 9 animals<br>74 sessions | Active DG Rest | 0.013 ± 0.002 | 9 animals<br>74 sessions | LMM | 4e-17 |
| 2a right A-A' | Passive V1 Run | 0.008 ± 0.002 | 4 animals<br>6 sessions | Passive V1 Rest | 0.020 ± 0.002 | 4 animals<br>6 sessions | LMM | 5e-17 |
| 2a right A-A' | Passive DG Run | 0.007 ± 0.002 | 4 animals<br>14 sessions | Passive DG Rest | 0.014 ± 0.002 | 4 animals<br>14 sessions | LMM | 3e-14 |
| 2b | A-B V1 Active | 0.023 ± 0.002 | 4 animals<br>10 sessions | A-B V1 Passive | 0.022 ± 0.002 | 4 animals<br>7 sessions | LMM | 0.347 |
| 2b | A-B DG Active | 0.011 ± 0.002 | 5 animals<br>30 sessions | A-B DG Passive | 0.011 ± 0.003 | 4 animals<br>14 sessions | LMM | 0.908 |
| 2b | A-A' V1 Active | 0.022 ± 0.002 | 4 animals<br>8 sessions | A-A' V1 Passive | 0.020 ± 0.002 | 4 animals<br>6 sessions | LMM | 0.644 |
| 2b | A-A' DG Active | 0.013 ± 0.002 | 9 animals<br>75 sessions | A-A' DG Passive | 0.014 ± 0.002 | 4 animals<br>14 sessions | LMM | 0.384 |
| **Figure 3** |  |  |  |  |  |  |  |  |
| 3d left active | Accuracy A-B V1 | 0.328 ± 0.020 | 4 animals<br>10 sessions | Zeroes | 0 | / | LMM | 6e-111 |
| 3d left active | Accuracy A-B DG | 0.070 ± 0.023 | 4 animals<br>30 sessions | Zeroes | 0 | / | LMM | 2e-6 |
| 3d left active | Accuracy A-A' V1 | 0.147 ± 0.028 | 4 animals<br>8 sessions | Zeroes | 0 | / | LMM | 2e-5 |
| 3d left active | Accuracy A-A' DG | 0.062 ± 0.023 | 9 animals<br>73 sessions | Zeroes | 0 | / | LMM | 1e-16 |
| 3d right passive | Accuracy A-B V1 | 0.175 ± 0.045 | 4 animals<br>7 sessions | Zeroes | 0 | / | LMM | 7e-4 |
| 3d right passive | Accuracy A-B DG | 0.013 ± 0.013 | 4 animals<br>14 sessions | Zeroes | 0 | / | LMM | 0.858 |
| 3d right passive | Accuracy A-A' V1 | 0.127 ± 0.049 | 4 animals<br>6 sessions | Zeroes | 0 | / | LMM | 1e-6 |
| 3d right passive | Accuracy A-A' DG | 0.020 ± 0.024 | 4 animals<br>14 sessions | Zeroes | 0 | / | LMM | 0.786 |
| 3e left V1 | Accuracy active A-B | 0.328 ± 0.020 | 4 animals<br>10 sessions | Accuracy active A-A' | 0.147 ± 0.028 | 4 animals<br>8 sessions | LMM | 1e-6 |
| 3e left V1 | Accuracy passive A-A' | 0.175 ± 0.045 | 4 animals<br>7 sessions | Accuracy passive A-A' | 0.127 ± 0.049 | 4 animals<br>6 sessions | LMM | 0.47 |
| 3e right DG | Accuracy active A-B | 0.070 ± 0.023 | 5 animals<br>30 sessions | Accuracy active A-A' | 0.062 ± 0.023 | 9 animals<br>73 sessions | LMM | 0.071 |
| 3e right DG | Accuracy passive A-A' | 0.013 ± 0.013 | 4 animals<br>14 sessions | Accuracy passive A-A' | 0.020 ± 0.024 | 4 animals<br>14 sessions | LMM | 0.99 |
| 3f left A-B | V1 active Slope | 0.008 ± 0.019 | 10 sessions | Accuracy function of running speed | / | / | LMM | 0.67 |
| 3f left A-B | V1 passive Slope | -0.005 ± 0.005 | 7 sessions | Accuracy function of running speed | / | / | LMM | 0.35 |
| 3f left A-B | DG active Slope | 0.005 ± 0.003 | 30 sessions | Accuracy function of running speed | / | / | LMM | 0.11 |
| 3f left A-B | DG passive Slope | -0.003 ± 0.042 | 14 sessions | Accuracy function of running speed | / | / | LMM | 0.94 |
| 3f right A-A' | V1 active Slope | -0.006 ± 0.007 | 8 sessions | Accuracy function of running speed | / | / | LMM | 0.34 |
| 3f right A-A' | V1 passive Slope | 0.003 ± 0.004 | 6 sessions | Accuracy function of running speed | / | / | LMM | 0.35 |
| 3f right A-A' | DG active Slope | 0.008 ± 0.001 | 73 sessions | Accuracy function of running speed | / | / | LMM | 2e-9 |
| 3f right A-A' | DG passive Slope | 0.002 ± 0.002 | 14 sessions | Accuracy function of running speed | / | / | LMM | 0.30 |
| 3g left A-B | V1 active Slope | -0.002 ± 0.005 | 10 sessions | Accuracy function of event rate | / | / | LMM | 0.74 |
| 3g left A-B | V1 passive Slope | 0.002 ± 0.003 | 7 sessions | Accuracy function of event rate | / | / | LMM | 0.55 |
| 3g left A-B | DG active Slope | 0.001 ± 0.046 | 30 sessions | Accuracy function of event rate | / | / | LMM | 0.99 |
| 3g left A-B | DG passive Slope | 0.002 ± 0.035 | 14 sessions | Accuracy function of event rate | / | / | LMM | 0.97 |
| 3g right A-A' | V1 active Slope | -0.002 ± 0.003 | 8 sessions | Accuracy function of event rate | / | / | LMM | 0.58 |
| 3g right A-A' | V1 passive Slope | -0.003 ± 0.271 | 6 sessions | Accuracy function of event rate | / | / | LMM | 0.99 |
| 3g right A-A' | DG active Slope | 0.001 ± 0.001 | 73 sessions | Accuracy function of event rate | / | / | LMM | 0.30 |
| 3g right A-A' | DG passive Slope | -0.001 ± 0.002 | 14 sessions | Accuracy function of event rate | / | / | LMM | 0.33 |
| 3h left A-B | V1 active Slope | -0.001 ± 0.002 | 4 animals<br>10 sessions | Accuracy function of neurons | / | / | LMM | 0.73 |
| 3h left A-B | V1 passive Slope | 0.005 ± 0.002 | 4 animals<br>7 sessions | Accuracy function of neurons | / | / | LMM | 0.0101 |
| 3h left A-B | DG active Slope | -0.000 ± 0.000 | 5 animals<br>30 sessions | Accuracy function of neurons | / | / | LMM | 0.38 |
| 3h right A-A' | V1 active Slope | -0.005 ± 0.010 | 4 animals<br>8 sessions | Accuracy function of neurons | / | / | LMM | 0.64 |
| 3h right<br>A-A' | V1 passive<br>*Slope* | $0.004 \pm 0.002$ | 4 animals<br>6 sessions | Accuracy function<br>of neurons | / | / | LMM | 0.08 |
| 3h right<br>A-A' | DG active<br>*Slope* | $-0.000 \pm 0.000$ | 9 animals<br>73 sessions | Accuracy function<br>of neurons | / | / | LMM | 0.82 |

