## Supplemental Figures and Table for "Task engagement differentially drives hippocampal and neocortical neural codes"

### Supplementary figures

*All statistical values are compiled in a dedicated table at the end of the supplementary figure section.*

#### Figure S1

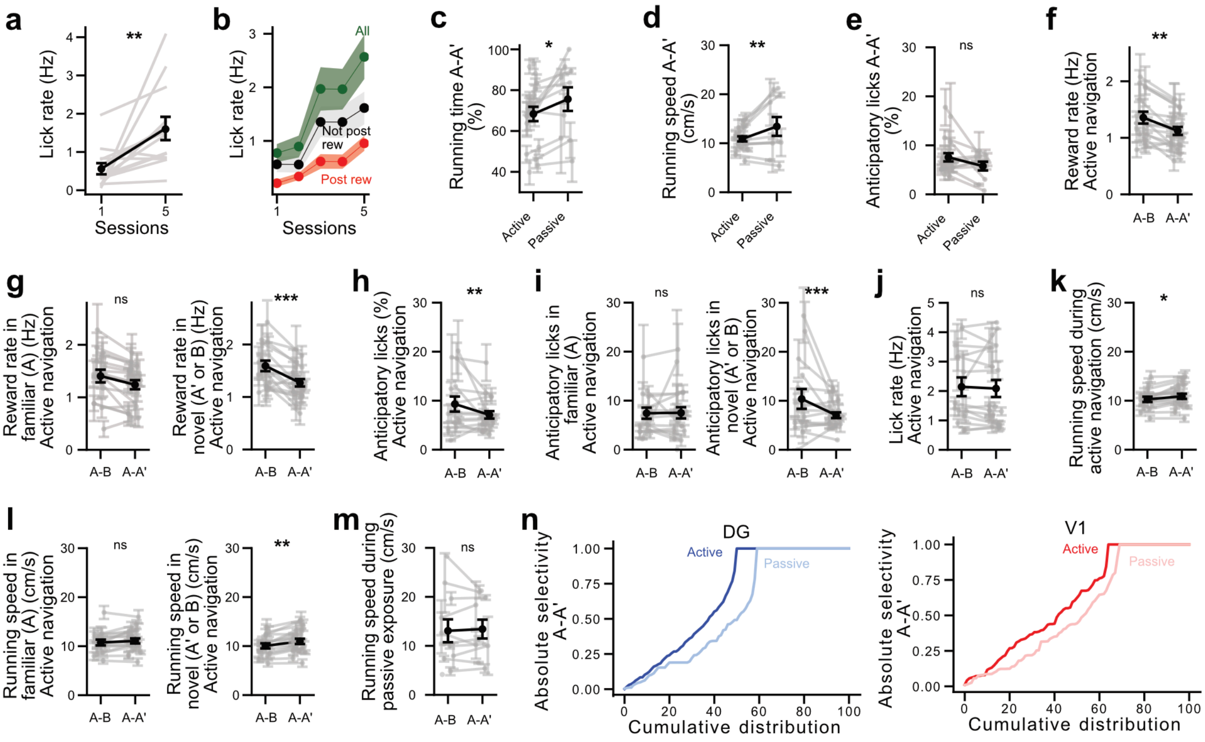

**Figure S1.** Supplementary to Fig. 1

**(a)** Comparison of lick rate between the first and fifth training sessions. **(b)** Lick rate increases with learning when considering all licks (green), licks following the delivery of a reward (‘consumption licks’, red) and non-consumption licks (black). **(c-e)** Comparison between active and passive conditions, in the environment pair A-A’, of **(c)** the running time, **(d)** the running speed and **(e)** the ratio of anticipatory licks. **(f)** Comparison of the reward rate during active navigation between A-B and A-A’. **(g)** Comparison of the reward rate within the familiar environment A depending on whether it is explored during an A-B or an A-A’ session (left), and between the two novel environments B and A’ (right). **(h)** and **(i)** are the same as respectively **(f)** and **(g)** for the ratio of anticipatory licks. **(j)** Comparison of the mean lick rate during active navigation between A-B and A-A’. **(k)** and **(l)** are the same as respectively **(f)** and **(g)** for the ratio of anticipatory licks. **(m)** Comparison of running speed during passive exposure within A-B and A-A’ environment pairs. **(n)** Distribution of absolute selectivity from individual neurons pulled across A-A’ recording sessions in active (darker colours) and passive (lighter) conditions, for DG (top) and V1 recordings (bottom).

In **(a)**, grey lines correspond to a single animal, while dark lines show the average across animals. Error bars indicate the standard error of the mean.

In **(b)**, solid lines show the mean while shaded areas correspond to the standard error of the mean.

In **(c-m)**, error bars indicate the standard error of the mean. Each grey line corresponds to multiple sessions from a single animal, while dark lines show the average across animals.

#### Figure S2

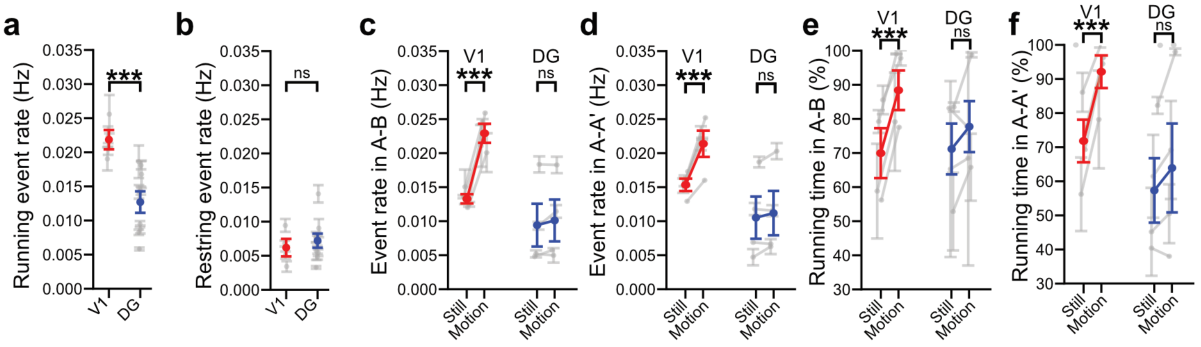


**Figure S2.** Supplementary to Fig. 2

**(a,b)** Comparison of the mean event rate during **(a)** running and **(b)** resting periods, both pairs of environments pulled together. **(c,d)** Comparison of the mean event rate during passive exposure between periods when the visual flow is still and periods when it is in motion for **(c)** the A-B pair of environments and **(d)** the A-A' pair. **(e,f)** Same as **(c,d)** but showing the ratio of time the animal spent running during the session.

In all panels, error bars indicate the standard error of the mean. Each grey line corresponds to multiple sessions from a single animal, while coloured lines show the average across animals.

### Figure S3

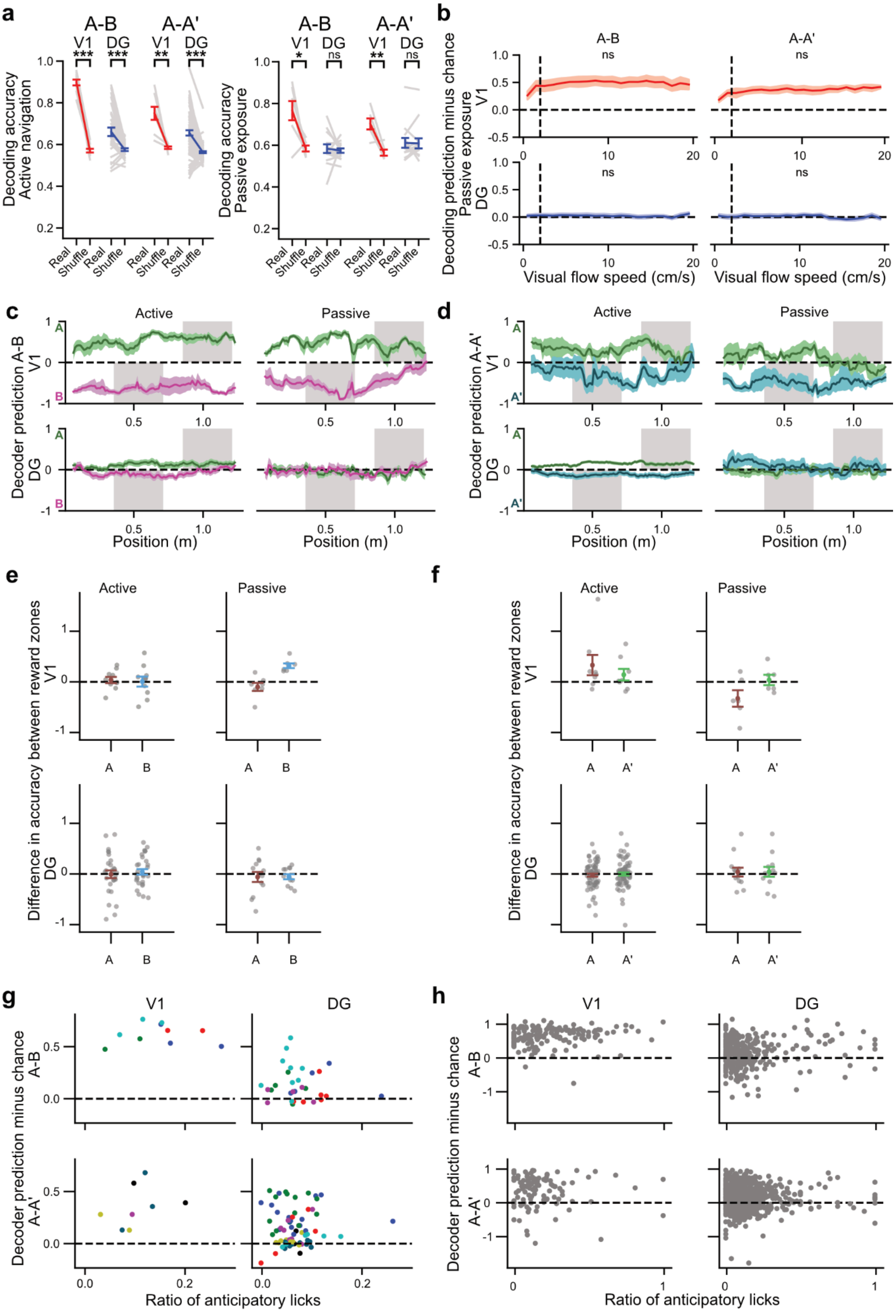

a
A-B
A-A'
V1
DG
Decoding accuracy
Active navigation
Decoding accuracy
Passive exposure
Real
Shuffle
b
Decoding prediction minus chance
Passive exposure
ns
Visual flow speed (cm/s)
c
d
Active
Passive
Decoder prediction A-B
Decoder prediction A-A'
Position (m)
e
f
Difference in accuracy between reward zones
g
h
Decoder prediction minus chance
Ratio of anticipatory licks

**Figure S3:** Supplementary to Fig. 3

**(a)** Comparison of raw decoding accuracy between real and shuffle data under active (left) and passive (right) conditions. **(b)** Decoding accuracy during passive exposure as a function of the visual flow speed (i.e. the motion within the VR movie, as opposed to running speed). **(c)** Decoding prediction as a function of position for the environment pair A-B. Prediction when visiting environment A (green) is positive when correct, whereas prediction when visiting environment B (magenta) is negative when correct. Gray shaded areas indicate reward zone location. **(d)** Same as (c) for the environment pair A-A'. The prediction when visiting environment A' (turquoise) is negative when correct. **(e)** Decoding accuracy within the reward zone boundaries minus decoding accuracy within another part of the track (taken between the limits of the reward zone from the opposite environment) for the environment A (explored during an A-B session) and B. **(f)** Same as (e) for the environment pair A-A'. **(g)** Decoding accuracy as a function of anticipatory licks ratio with session as a statistical unit. **(h)** Same as (g) with laps as a statistical unit.

In **(a)**, error bars indicate the standard error of the mean. Each grey line corresponds to multiple sessions from a single animal, while coloured lines show the average across animals.

In **(b-d)**, solid lines show the mean while shaded areas correspond to the standard error of the mean.

In **(e,f)**, error bars indicate the standard error of the mean. Each dot is a single session.

In **(g)**, each dot corresponds to a single session, colours represent animals.

In **(h),** each dot is a single lap.

#### Figure S4

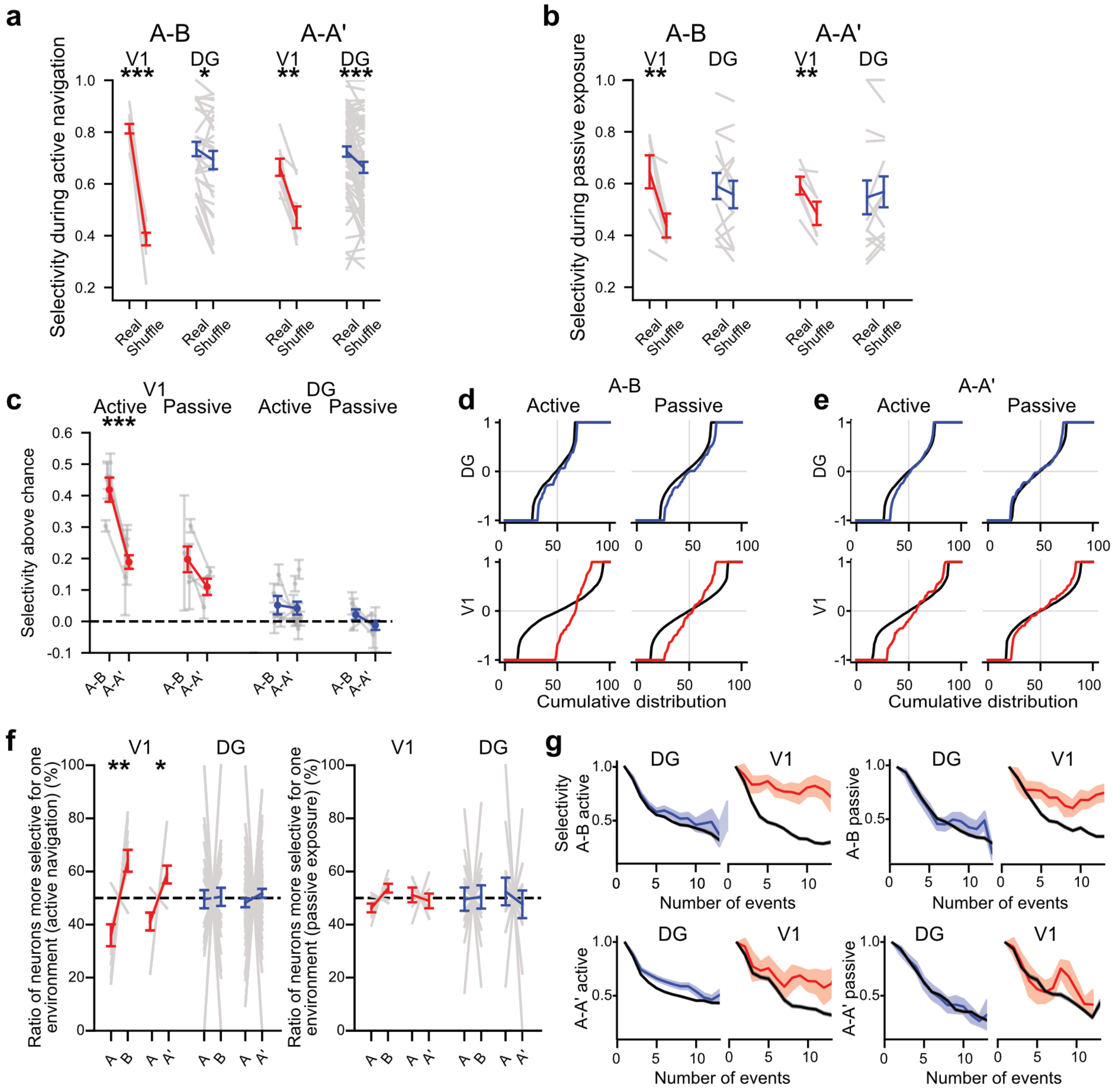


**Figure S4:** During passive exposure, selectivity of neuronal activity drops to chance levels in the dentate gyrus but not in V1.

**(a)** Comparison between real and chance selectivity during active navigation. **(b)** Comparison between real and chance selectivity during passive exposure. **(c)** Comparison of normalised selectivity between pairs of environments. **(d)** Signed selectivity distributions from real (coloured) and shuffled (black) data in the A-B environment. **(d)** Same as **(d)** for the A-A' pair of environments. **(f)** Proportion of neurons preferentially firing in one or another environment during active navigation (left) and passive exposure (right). **(g)** Selectivity as a function of the number of events, for real (coloured) and shuffled (black) data.

In **(a,b),** error bars indicate the standard error of the mean. Grey lines correspond to a single session, while coloured lines show the average across animals.

In **(c,f)**, error bars indicate the standard error of the mean. Grey lines correspond to multiple sessions from a single animal, while coloured lines show the average across animals.

In **(g)**, solid lines show the mean while shaded areas correspond to the standard error of the mean.

#### Figure S5

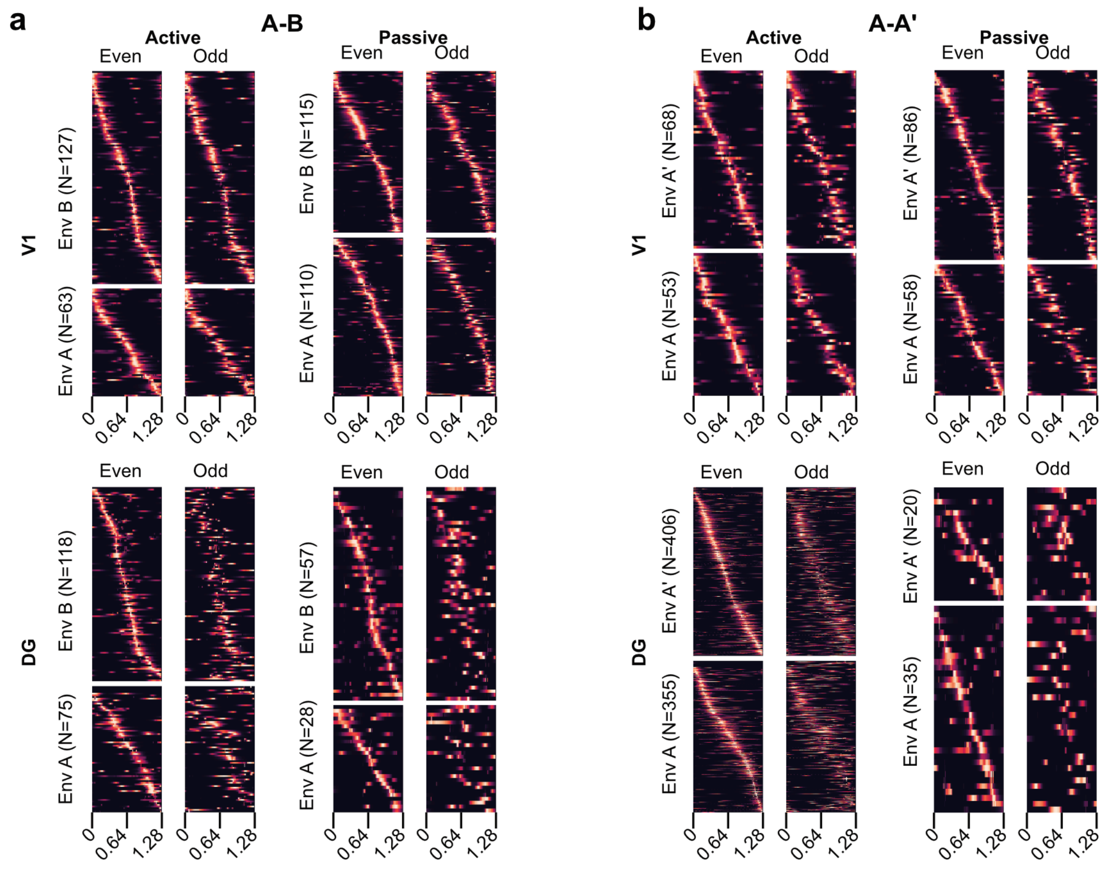


**Figure S5.** Cells with consistent place fields can be identified in all regions and conditions

Cross-validated heatmaps for identified spatially modulated cells across all conditions. Laps are divided into two halves (odd/even), and neurons are sorted based on the location of their peak event rate within even laps. This sorting is conserved when displaying odd laps. Each neuron event rate is normalised to its peak activity. Lighter colours indicate a higher event rate. **(a)** A-B pair of environments. **(b)** A-A' pair.

#### Statistics (supplementary figures)

*LMM: Linear Mixed Model; TT: paired t-test; MW: Mann–Whitney U test; WIL: Wilcoxon signed-rank test; BENF: Benferroni correction for multiple comparisons.*

*When providing N values for linear mixed models, the first row corresponds to the number of groups and the second row to the number of observations.*

| Panel | Field 1 | Value | N | Field 2 | Value | N | Test | p-val |
|---|---|---|---|---|---|---|---|---|
| **Figure S1** | | | | | | | | |
| S1a | Lick rate session 1 | 0.56 ± 0.15 | 13 animals | Lick rate session 5 | 1.61 ± 0.30 | 13 animals *(paired)* | TT | 0.002 |
| S1c | Running time A-A' active | 68.3 ± 3.5 | 20 animals 260 sessions | Running time A-A' passive | 75.6 ± 5.8 | 10 animals 28 sessions | LMM | 0.038 |
| S1d | Running speed A-A' active | 10.9 ± 0.5 | 20 animals 260 sessions | Running speed A-A' passive | 13.4 ± 1.9 | 10 animals 28 sessions | LMM | 0.002 |
| S1e | Anticipatory licks A-A' active | 7.6 ± 0.8 | 20 animals 252 sessions | Anticipatory licks A-A' passive | 5.8 ± 0.9 | 10 animals 28 sessions | LMM | 0.514 |
| S1f | Reward rate active A-B | 1.36 ± 0.11 | 16 animals 145 sessions | Reward rate active A-A' | 1.12 ± 0.07 | 16 animals 260 sessions | LMM | 0.006 |
| S1g left | Reward rate in familiar active A-B | 1.40 ± 0.12 | 16 animals 143 sessions | Reward rate in familiar active A-A' | 1.24 ± 0.09 | 20 animals 258 sessions | LMM | 0.147 |
| S1g right | Reward rate in novel active A-B | 1.59 ± 0.10 | 16 animals 143 sessions | Reward rate in novel active A-A' | 1.27 ± 0.07 | 20 animals 259 sessions | LMM | 0.001 |
| S1h | Anticipatory licks active A-B | 9.3 ± 1.5 | 15 animals 98 sessions | Anticipatory licks active A-A' | 7.1 ± 0.8 | 20 animals 201 sessions | LMM | 0.007 |
| S1i left | Anticipatory licks in familiar active A-B | 7.4 ± 1.1 | 15 animals 95 sessions | Anticipatory licks in familiar active A-A' | 7.5 ± 1.1 | 20 animals 197 sessions | LMM | 0.534 |
| S1i right | Anticipatory licks in novel active A-B | 10.4 ± 2.0 | 15 animals 97 sessions | Anticipatory licks in novel active A-A' | 7.0 ± 0.6 | 20 animals 199 sessions | LMM | 3.8e-4 |
| S1j | Lick rate active A-B | 2.14 ± 0.32 | 15 animals 98 sessions | Lick rate active A-A' | 2.08 ± 0.29 | 20 animals 201 sessions | LMM | 0.37 |
| S1k | Running speed active A-B | 10.3 ± 0.5 | 16 animals 145 sessions | Running speed active A-A' | 10.9 ± 0.5 | 20 animals 260 sessions | LMM | 0.018 |
| S1l left | Running speed in familiar active A-B | 10.7 ± 0.6 | 16 animals 143 sessions | Running speed in familiar active A-A' | 11.1 ± 0.5 | 20 animals 258 sessions | LMM | 0.117 |
| S1l right | Running speed in novel active A-B | 10.0 ± 0.5 | 16 animals 143 sessions | Running speed in novel active A-A' | 11.0 ± 0.5 | 20 animals 259 sessions | LMM | 0.004 |
| S1m | Running speed passive A-B | 13.1 ± 2.3 | 11 animals 28 sessions | Running speed passive A-A' | 13.4 ± 1.9 | 10 animals 28 sessions | LMM | 0.985 |
| **Figure S2** | | | | | | | | |
| S2a | Running event rate V1 | 0.022 ± 0.001 | 4 animals 31 sessions | Running event rate DG | 0.013 ± 0.002 | 9 animals 133 sessions | MW | 2.7e-12 |
| S2b | Resting event rate V1 | 0.006 ± 0.001 | 4 animals 31 sessions | Resting event rate DG | 0.007 ± 0.001 | 9 animals 133 sessions | MW | 0.89 |
| S2c | Event rate V1 still | 0.013 ± 0.001 | 4 animals 7 sessions | Event rate V1 motion | 0.023 ± 0.001 | 4 animals 7 sessions | LMM | 2e-10 |
| S2c | Event rate DG still | 0.009 ± 0.003 | 4 animals 14 sessions | Event rate DG motion | 0.010 ± 0.003 | 4 animals 14 sessions | LMM | 0.26 |
| S2d | Event rate V1 still | 0.015 ± 0.001 | 4 animals 6 sessions | Event rate V1 motion | 0.021 ± 0.002 | 4 animals 6 sessions | LMM | 5e-12 |
| S2d | Event rate DG still | 0.011 ± 0.003 | 4 animals 14 sessions | Event rate DG motion | 0.011 ± 0.003 | 4 animals 14 sessions | LMM | 0.29 |

| | | | | | | | | |
|---|---|---|---|---|---|---|---|---|
| S2e | Running time V1 still | 70.0 ± 7.3 | 4 animals 7 sessions | Running time V1 motion | 88.4 ± 5.8 | 4 animals 7 sessions | LMM | 8e-4 |
| S2e | Running time DG still | 71.2 ± 7.4 | 4 animals 14 sessions | Running time DG motion | 77.7 ± 7.5 | 4 animals 14 sessions | LMM | 0.16 |
| S2f | Running time V1 still | 71.9 ± 6.3 | 4 animals 6 sessions | Running time V1 motion | 92.3 ± 4.8 | 4 animals 6 sessions | LMM | 0.001 |
| S2f | Running time DG still | 57.5 ± 9.5 | 4 animals 14 sessions | Running time DG motion | 64.0 ± 13.0 | 4 animals 14 sessions | LMM | 0.21 |
| **Figure S3** | | | | | | | | |
| S3a left active | A-B V1 real | 0.897 ± 0.015 | 10 sessions | A-B V1 shuffle | 0.572 ± 0.007 | 10 sessions *(paired)* | TT | 6.1e-9 |
| S3a left active | A-B DG real | 0.659 ± 0.019 | 30 sessions | A-B DG shuffle | 0.580 ± 0.006 | 30 sessions (paired) | TT | 1e-4 |
| S3a left active | A-A' V1 real | 0.742 ± 0.033 | 8 sessions | A-A' V1 shuffle | 0.586 ± 0.006 | 8 sessions (paired) | TT | 0.003 |
| S3a left active | A-A' DG real | 0.653 ± 0.012 | 73 sessions | A-A' DG shuffle | 0.565 ± 0.005 | 73 sessions (paired) | TT | 2e-10 |
| S3a right passive | A-B V1 real | 0.766 ± 0.046 | 7 sessions | A-B V1 shuffle | 0.583 ± 0.014 | 7 sessions (paired) | TT | 0.014 |
| S3a right passive | A-B DG real | 0.586 ± 0.019 | 14 sessions | A-B DG shuffle | 0.582 ± 0.009 | 14 sessions (paired) | TT | 0.86 |
| S3a right passive | A-A' V1 real | 0.703 ± 0.026 | 6 sessions | A-A' V1 shuffle | 0.564 ± 0.015 | 6 sessions (paired) | TT | 0.008 |
| S3a right passive | A-A' DG real | 0.613 ± 0.025 | 14 sessions | A-A' DG shuffle | 0.609 ± 0.024 | 14 sessions (paired) | TT | 0.79 |
| S3b | V1 A-B *Slope* | 0.000 ± 0.001 | 7 sessions | Accuracy function of visual flow | / | / | LMM | 0.88 |
| S3b | V1 A-A' *Slope* | 0.003 ± 0.024 | 6 sessions | Accuracy function of visual flow | / | / | LMM | 0.91 |
| S3b | DG A-B *Slope* | -0.001 ± 0.001 | 14 sessions | Accuracy function of visual flow | / | / | LMM | 0.49 |
| S3b | DG A-A' *Slope* | -0.001 ± 0.044 | 14 sessions | Accuracy function of visual flow | / | / | LMM | 0.99 |
| S3e | V1 active A | 0.04 ± 0.06 | 10 sessions | Zeroes | 0 | / | WIL + BENF | 1.00 |
| S3e | V1 active B | 0.01 ± 0.10 | 10 sessions | Zeroes | 0 | / | WIL + BENF | 1.00 |
| S3e | V1 passive A | -0.10 ± 0.08 | 7 sessions | Zeroes | 0 | / | WIL + BENF | 1.00 |
| S3e | V1 passive B | 0.32 ± 0.05 | 7 sessions | Zeroes | 0 | / | WIL + BENF | 0.31 |
| S3e | DG active A | -0.01 ± 0.08 | 29 sessions | Zeroes | 0 | / | WIL + BENF | 1.00 |
| S3e | DG active B | 0.04 ± 0.05 | 30 sessions | Zeroes | 0 | / | WIL + BENF | 1.00 |
| S3e | DG passive A | -0.06 ± 0.10 | 13 sessions | Zeroes | 0 | / | WIL + BENF | 1.00 |
| S3e | DG passive B | -0.06 ± 0.05 | 12 sessions | Zeroes | 0 | / | WIL + BENF | 1.00 |
| S3f | V1 active A | 0.33 ± 0.20 | 8 sessions | Zeroes | 0 | / | WIL + BENF | 0.85 |
| S3f | V1 active A' | 0.14 ± 0.11 | 8 sessions | Zeroes | 0 | / | WIL + BENF | 1.00 |

| | | | | | | | | |
|---|---|---|---|---|---|---|---|---|
| S3f | V1 passive<br>A | -0.33 ± 0.16 | 6 sessions | Zeroes | 0 | / | WIL +<br>BENF | 0.97 |
| S3f | V1 passive<br>A' | 0.04 ± 0.10 | 6 sessions | Zeroes | 0 | / | WIL +<br>BENF | 1.00 |
| S3f | DG active<br>A | -0.02 ± 0.03 | 71 sessions | Zeroes | 0 | / | WIL +<br>BENF | 1.00 |
| S3f | DG active<br>A' | -0.00 ± 0.03 | 73 sessions | Zeroes | 0 | / | WIL +<br>BENF | 1.00 |
| S3f | DG passive<br>A | 0.04 ± 0.09 | 13 sessions | Zeroes | 0 | / | WIL +<br>BENF | 1.00 |
| S3f | DG passive<br>A' | 0.04 ± 0.10 | 12 sessions | Zeroes | 0 | / | WIL +<br>BENF | 1.00 |
| S3g | V1 A-B<br>*Slope* | 0.030 ± 0.519 | 4 animals<br>10 sessions | Accuracy function<br>of anticipatory licks | / | / | LMM | 0.95 |
| S3g | DG A-B<br>*Slope* | -0.041 ± 0.659 | 5 animals<br>30 sessions | Accuracy function<br>of anticipatory licks | / | / | LMM | 0.95 |
| S3g | V1 A-A'<br>*Slope* | 1.384 ± 1.13 | 4 animals<br>8 sessions | Accuracy function<br>of anticipatory licks | / | / | LMM | 0.22 |
| S3g | DG A-A'<br>*Slope* | 0.227 ± 0.125 | 9 animals<br>73 session | Accuracy function<br>of anticipatory licks | / | / | LMM | 0.069 |
| S3h | V1 A-B<br>*Slope* | 0.195 ± 0.121 | 10 sessions<br>161 laps | Accuracy function<br>of anticipatory licks | / | / | LMM | 0.10 |
| S3h | DG A-B<br>*Slope* | 0.268 ± 0.171 | 29 sessions<br>304 laps | Accuracy function<br>of anticipatory licks | / | / | LMM | 0.12 |
| S3h | V1 A-A'<br>*Slope* | -0.234 ± 0.538 | 8 sessions<br>108 laps | Accuracy function<br>of anticipatory licks | / | / | LMM | 0.66 |
| S3h | DG A-A'<br>*Slope* | -0.047 ± 0.088 | 72 sessions<br>722 laps | Accuracy function<br>of anticipatory licks | / | / | LMM | 0.59 |
| **Figure S4** | | | | | | | | |
| S4a | A-B<br>V1 real | 0.813 ± 0.018 | 10 sessions | A-B<br>V1 shuffle | 0.387 ± 0.024 | 10 sessions<br>*(paired)* | TT | 3.6e-7 |
| S4a | A-B<br>DG real | 0.735 ± 0.028 | 30 sessions | A-B<br>DG shuffle | 0.692 ± 0.035 | 30 sessions<br>*(paired)* | TT | 0.014 |
| S4a | A-A'<br>V1 real | 0.664 ± 0.033 | 8 sessions | A-A'<br>V1 shuffle | 0.471 ± 0.042 | 8 sessions<br>*(paired)* | TT | 0.002 |
| S4a | A-A'<br>DG real | 0.725 ± 0.020 | 74 sessions | A-A'<br>DG shuffle | 0.664 ± 0.021 | 74 sessions<br>*(paired)* | TT | 1.2e-6 |
| S4b | A-B<br>V1 real | 0.645 ± 0.064 | 7 sessions | A-B<br>V1 shuffle | 0.438 ± 0.046 | 7 sessions<br>*(paired)* | TT | 0.008 |
| S4b | A-B<br>DG real | 0.590 ± 0.050 | 14 sessions | A-B<br>DG shuffle | 0.558 ± 0.053 | 14 sessions<br>*(paired)* | TT | 0.20 |
| S4b | A-A'<br>V1 real | 0.592 ± 0.034 | 6 sessions | A-A'<br>V1 shuffle | 0.485 ± 0.045 | 6 sessions<br>*(paired)* | TT | 0.007 |
| S4b | A-A'<br>DG real | 0.547 ± 0.065 | 14 sessions | A-A'<br>DG shuffle | 0.568 ± 0.060 | 14 sessions<br>*(paired)* | TT | 0.34 |
| S4c | V1<br>active A-B | 0.42 ± 0.04 | 4 animals<br>10 sessions | V1<br>active A-A' | 0.19 ± 0.02 | 4 animals<br>8 sessions | LMM | 5.9e-6 |
| S4c | V1<br>passive A-B | 0.20 ± 0.04 | 4 animals<br>7 sessions | V1<br>passive A-A' | 0.11 ± 0.03 | 4 animals<br>6 sessions | LMM | 0.11 |
| S4c | DG<br>active A-B | 0.05 ± 0.03 | 5 animals<br>30 sessions | DG<br>active A-A' | 0.04 ± 0.02 | 9 animals<br>74 sessions | LMM | 0.28 |
| S4c | DG<br>passive A-B | 0.02 ± 0.02 | 4 animals<br>14 sessions | DG<br>passive A-A' | -0.01 ± 0.01 | 4 animals<br>14 sessions | LMM | 0.10 |

| | | | | | | | | |
|---|---|---|---|---|---|---|---|---|
| S4f left (active) | V1 A vs B A | 36.0 ± 4.1 | 10 sessions | V1 A vs B B | 64.0 ± 4.1 *(1 - field 1)* | 10 sessions *(paired)* | TT | 0.008 |
| S4f left (active) | V1 A vs A' A | 41.2 ± 3.4 | 8 sessions | V1 A vs A' A' | 58.8 ± 3.4 *(1 - field 1)* | 8 sessions *(paired)* | TT | 0.03 |
| S4f left (active) | DG A vs B A | 49.6 ± 3.4 | 30 sessions | DG A vs B B | 50.4 ± 3.4 *(1 - field 1)* | 30 sessions *(paired)* | TT | 0.90 |
| S4f left (active) | DG A vs A' A | 48.2 ± 1.7 | 74 sessions | DG A vs A' A' | 51.8 ± 1.7 *(1 - field 1)* | 74 sessions *(paired)* | TT | 0.28 |
| S4f right (passive) | V1 A vs B A | 46.3 ± 1.6 | 7 sessions | V1 A vs B B | 53.7 ± 1.6 *(1 - field 1)* | 7 sessions *(paired)* | TT | 0.06 |
| S4f right (passive) | V1 A vs A' A | 51.1 ± 2.8 | 6 sessions | V1 A vs A' A' | 48.9 ± 2.8 *(1 - field 1)* | 6 sessions *(paired)* | TT | 0.70 |
| S4f right (passive) | DG A vs B A | 49.6 ± 4.4 | 14 sessions | DG A vs B B | 50.4 ± 4.4 *(1 - field 1)* | 14 sessions *(paired)* | TT | 0.93 |
| S4f right (passive) | DG A vs A' A | 52.4 ± 5.2 | 14 sessions | DG A vs A' A' | 47.6 ± 5.2 *(1 - field 1)* | 14 sessions *(paired)* | TT | 0.65 |
